# Transcriptional profiling of human brain cortex identifies novel lncRNA-mediated networks dysregulated in amyotrophic lateral sclerosis

**DOI:** 10.1101/2024.03.18.585481

**Authors:** Alessandro Palma, Monica Ballarino

## Abstract

Amyotrophic lateral sclerosis (ALS) is a neurodegenerative disease for which a comprehensive knowledge about the pathological mechanisms is still lacking. A multitude of dysregulated cellular processes and pathways have been linked to ALS so far, including the recent focus directed toward the implication of several classes of non-coding (nc)RNAs. Within this context, the class of long ncRNAs (lncRNAs), may provide an important contribution to the onset and the severity of ALS pathogenesis, due to their high tissue specificity and their function as gene expression regulators. Nevertheless, their identification in humans often relies on differential expression analyses from bulk RNA-seq, which limits their targeting in the cellular contexts where they may be primarily involved.

Here we apply dedicated pipelines to single-nucleus nuclei datasets to study lncRNA from non-pathological and pre-frontal ALS human cortex. We found that in brain, distinct cell subtypes express very different pattern of lncRNAs to suggest possible roles in cellular processes found dysregulated in ALS patients. Moreover, we show the lncRNA involvement in important gene regulatory networks that result differentially regulated in pathological conditions and dissect the genomic organization of differentially expressed lncRNAs.

## Introduction

Amyotrophic Lateral Sclerosis (ALS) represents one of the most common neurodegenerative disorders, characterized by the progressive loss of upper and lower motor neurons in the brain cortex, brainstem and spinal cord ^1^, with a worldwide incidence of ∼2 per 100,000 person-years ^2^. From cellular and molecular perspectives, ALS is associated with protein aggregation and cellular stress, as well as inflammation and oxidative damage, ultimately leading to muscle weakness and death of motor neurons. The accumulation of misfolded proteins, such as superoxide dismutase 1 (SOD1) ^3^, in motor neurons can alter normal cellular processes and lead to neuronal damage. Additionally, mutations in genes such as chromosome 9 open reading frame 72 (C9orf72)^4,5^, transactive response DNA binding protein (TARDBP, coding for TDP-43 protein) ^6^, and fused in sarcoma (FUS) ^7^ have been identified in some cases of sporadic and familial ALS, highlighting, at least in part, the genetic component of the disease. Several other genes have been linked to ALS so far, including autophagy-related genes (SQSTM1, OPTN, C9orf72, TBK1) ^8–11^, altered ribostasis (HNRNPA2B1, HNRNPA1, MATR3), DNA damage ^12^, among others (reviewed in ^13^).

The involvement of RNA processing, metabolism and regulation in ALS pathogenesis has been described by numerous reports ^14–20^, and the importance of ribostasis - the appropriate production and regulation of the cellular transcriptome - is further confirmed by the fact that many neuromuscular and neurodegenerative disorders are associated to alterations in protein folding, RNA regulation, RNA binding proteins, or RNP aggregates ^17^. Such a scenario implies that distinct, yet uncovered, “molecular regulators” could be involved in ALS pathogenesis, including long non-coding RNAs (lncRNAs) ^21,22^ a novel class of transcripts whose main function relies on the regulation of biological functions related to cell differentiation and regeneration ^23–26^. By definition, they are non-coding transcripts longer than 500 nucleotides, mostly generated by the RNA-Pol II ^27^. Their physiological abilities rely on a wide-range of transcriptional and post-transcriptional mechanisms and on their peculiar subcellular localization, being able to interact with DNA, RNA and proteins in both the cytoplasm and nucleus ^28–32^. Several tools for studying lncRNAs have been developed so far ^33^, but a comprehensive understanding of their role and involvement in human diseases is still lacking.

Despite a very few reports mentioned a role or a function directly attributed to lncRNAs in this disease ^21,22,34,35^, some authors recently reported a differential expression of lncRNAs in ALS transcriptomic data. For instance, RNA sequencing performed on cervical, thoracic and lumbar regions from post-mortem ALS patients, displayed a statistically significant down-regulation of lncRNAs in ALS patients, accounting for almost 55% of all the down-regulated genes and 15% of the up-regulated ones ^36^. Many other transcriptomics data include lncRNAs among differentially expressed transcripts, which remained ignored due to the lack of functional annotation and the possibility to link their expression to the de-regulation of specific pathways and cellular processes.

As genetics studies have only partially dissected ALS pathogenesis, which remains limited to a small percentage of patients bearing the already identified driver mutations, understanding the complex cellular and molecular mechanisms of gene regulation in ALS in a “non-coding RNA” perspective is critical for the development of effective treatments for this neurodegenerative disorder, for which we urge for advancements.

Here we dissect the transcriptomic landscape of ALS using a publicly available single-nuclei RNA sequencing (snRNAseq) datasets from ALS and pathologically normal (PN) patients ^37^, and applied functional genomics approaches to search for cellular and transcriptional mechanisms that were dysregulated in ALS condition. We found several brain-expressed and brain-specific lncRNAs with potential roles in contributing to ALS dysregulated mechanisms occurring within neuronal cell populations.

## Results

### LncRNAs as biomarkers of distinct brain cell populations

To identify novel cell-type specific lncRNAs, we employed publicly available single-nucleus RNAseq datasets of brain cortex from 23 ALS patients and 17 healthy controls ^37^. After quality filtering for mitochondrial content and transcript number (see Materials and Methods), clustering analysis led to the identification of 27 clusters (Supplementary Fig. 1a) representative of distinct cell populations that were collectively regrouped into 7 major cell types, according to the expression of distinct markers and as already annotated by the authors of the original paper: Excitatory Neurons, Inhibitory Neurons, Oligodendrocytes, OPC (Oligodendrocyte Progenitor Cells), Microglia, Astrocytes and Vascular cells (Fig. 1a, b).

**Fig. 1.**
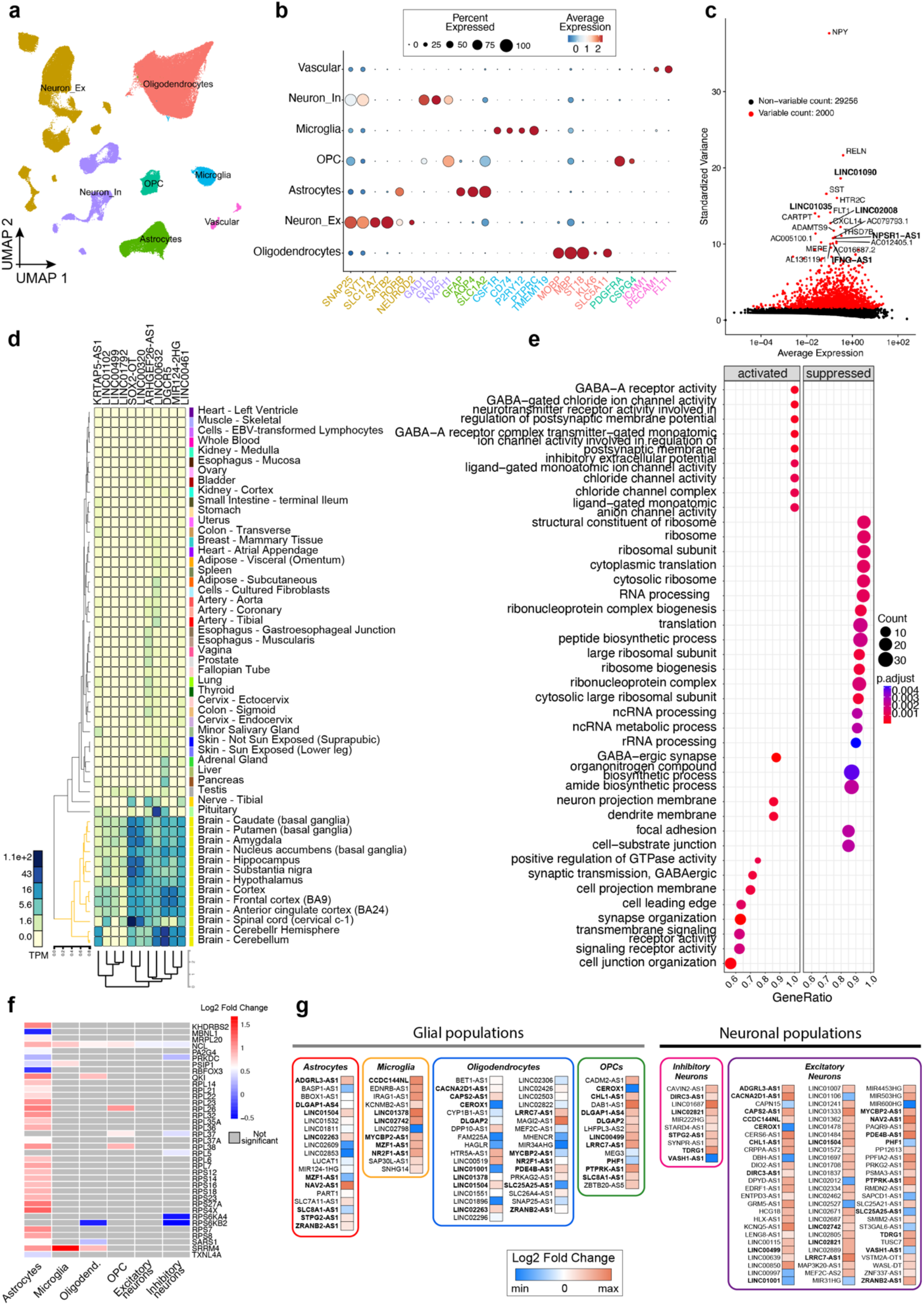
Single nuclei RNA-seq analysis of cell populations and lncRNAs. (a) UMAP projection of the identified populations, Oligodendrocytes, Neuron Excitatory (-Ex), Neuron Inhibitory (-In), Vascular, Oligodendrocyte Progenitor Cells (OPC), Microglia and Astrocytes. (b) Dotplot showing the type and the normalized expression of the readout markers used for cells identification. For each population, the average and the percentage of expression are reported. (c) Dotplot showing variable (red) and non-variable features (black) of the cell populations. LncRNAs are represented in bold. (d) Heatmap from GTEx resource showing the expression levels (TPM) of selected lncRNAs in the different tissues. (e) Dotplot of GSEA analysis of DGE astrocytes, showing significantly activated and suppressed processes among the top 20 GO Biological Process, Cell Component and Molecular Function. (f) Heatmap of a selected list of ribosomal-related genes showing their log2 fold changes in the different cell populations (ALS vs PN). (g) Heatmap of lncRNAs log2 fold change (ALS vs PN) for each cell population. LncRNAs that are shared between two or more populations are highlighted in bold.

Among the top 2000 variable genes, we found 286 transcripts (14.3%) classified as lncRNAs according to GENECODE release 44 (GRCh38.p14) (Fig. 1c). Moreover, markers computed for each cell population revealed the existence of 386 lncRNAs in total (Supplementary Table 4), the majority of which belonging to excitatory neurons and vascular cells (Supplementary Fig. 1b), and with some of them (20.9%) being strongly up- or down-regulated (|log2FC| > 2). Due to the known tissue specificity of lncRNAs, we queried GTEx database ^38^ for their expression in human tissues. Among the 386 lncRNAs, we found 11 lncRNAs almost uniquely present in neuronal and non-neuronal populations residing in the brain (Fig. 1d), six of which are expressed in brain vascular cells (LINC00632, SOX2-OT, LINC00320, DGCR5, MIR124-2HG and LINC00461), two in astrocytes (ARHGEF26-AS1 and LINC000499), one in oligodendrocytes (LINC01792), and two (LINC01102 and KRTAP5-AS1) in inhibitory and excitatory neurons, respectively.

Overall, these results suggest these species as general cell populations rather than ALS biomarkers (Supplementary Table 1), when merely considering differential gene expression testing among the distinct neuronal cell populations.

### LncRNAs may regulate distinct cellular processes in ALS glia

Single-cell RNA-seq datasets give the unique opportunity to study tissues at cell population level. This is crucial for the analysis of complex diseases like ALS, in which several cell populations could play different roles in disease pathogenesis. To date, many reports relied on the use of bulk transcriptomic, which hampers the possibility of assigning a specific cell type to any given dysregulated process. To fill this gap and identify ALS-linked lncRNAs in any given cell type, we applied pseudobulk analysis to ALS and PN samples across the different cell subtypes and found, for each comparison, interesting lncRNA signatures. Together with the previous observation that many lncRNAs are among the most variable features within the dataset, and the high percentage of lncRNA biomarkers, this fact prompted us to further investigate their possible role in ALS pathogenesis.

In terms of cell number, we could not find significant differences when comparing ALS to PN although we observed a decreased number of astrocytes in ALS patients (Supplementary Fig. 1c). After performing a pseudobulk analysis on astrocytes population (ALS *vs* PN), the Gene Set Enrichment Analysis (GSEA) of GO Biological Processes, Cellular Components and Molecular Functions performed on DEGs (adjusted p-value < 0.05) (Supplementary Table 1) highlighted the suppression of RNA-related processes, including RNA processing, ribonucleoprotein complex biogenesis, ribonucleoprotein complex, ribosome biogenesis, as well as non-coding RNA-related processes (ncRNA processing, ncRNA metabolic process) (Fig. 1 e). RNA-linked processes seemed to be particularly enriched in astrocytes with no detected de-regulated ribosomal genes in vascular population, and with only a few ribosomal genes in the other neuronal and glial cells (Fig. 1f). Other processes resulted activated, such as GABA-ergic signaling-related terms and synaptic organization and transmission, indicating a possible hyperactivation of synaptic transmission in this cell population. These GO terms were among the top 20 enriched terms in GSEA (Supplementary Table 2), which highlighted a possible alteration of RNA regulation in ALS astrocytes. Although the molecular mechanisms leading to astrogliosis are still far from being well characterized, this is in line with the impact of glial composition on disease progression ^36^, and the neurotoxic effect of astrocytes towards neuronal cell populations previously reported in rodent models of ALS ^39–41^. We also noted that 19 out of 332 differentially expressed transcripts were lncRNAs (Fig. 1g), 15 of which were up-regulated with a log2 fold change > 1. The majority of these lncRNAs were annotated as anti-sense transcripts, with their sense counterparts involved in splicing regulation (ZRANB2), ion transport (SLC8A1, SLC7A11), neuron guidance and development (ADGRL3, NAV2), post-synaptic density (DLGAP1), and transcriptional regulation (MZF1), according to Gene Ontology. Importantly, BASP1-AS1 was the only anti-sense transcript found up-regulated in concomitance to its sense counterpart. To note, the expression of BASP1 was previously found to be altered in ALS frontal cortex ^42^, while its anti-sense transcript was predicted by catRAPID ^43^ to bind SAFB2 (Supplementary Fig. 1d), an RNA-binding protein interacting with GGGGCC repeats ^44^, which are known to be enriched in C9orf72-mutated ALS patients. These results suggest an important impairment of RNA regulation in ALS astrocytes and a putative involvement of lncRNAs in these detrimental processes occurring in ALS.

Pseudobulk analysis performed on ALS *vs* PN microglial population led us to identify 363 DEGs, the majority of which (92.3%) were up-regulated in ALS samples. Further GSEA analysis revealed that several GO terms related to chromatin, DNA packaging and nucleosome organization resulted enriched in ALS patients (Supplementary Table 2). Furthermore, we observed the up-regulation of anti-sense RNAs whose sense counterparts act as DNA-binding transcription factors, such as NR2F1-AS1, MZF1-AS1 and MYCBP2-AS1. Other differentially expressed lncRNAs could be involved in the transcriptional regulation of genes directly involved in processes related to calcium signaling, including IRAG1-AS1 (antisense to the regulator of IP3-induced calcium release, IRAG1), EDNRB-AS1 (antisense to the activator of the phosphatidylinositol-calcium second messenger system, EDNRB) and KCNMB2-AS1 (antisense to the calcium-sensitive potassium channel, KCNMB2). Neurodegeneration, Amyotrophic Lateral Sclerosis, and other neurodegenerative disorders, were among the top 10 enriched KEGG pathways (Supplementary Table 3), as well as oxidative stress, recognized as a hallmark of ALS ^45^.

Oligodendrocytes represent the largest glial population in both ALS and PN patients, with a mean of 42.25% in ALS and 47.6% in PN samples. We could not find any significant change in oligodendrocytes or Oligodendroyte progenitor cell (OPC) numbers between ALS and PN, which is in line with other reports showing that an increase in oligodendrocytes proliferation may be accompanied by cell loss ^46^, the latter being hard to detect by snRNAseq. Differential expression analysis between ALS and PN samples, led to the identification of 903 statistically significant transcripts (adj. p-value < 0.05), 699 of which were up-regulated and 204 down-regulated in ALS. DEGs were enriched for suppressed processes related to RNA functions, including ribosomal and tRNA processing, non-coding RNA and RNA modification, as well as mitochondrial function (Supplementary Table 2). Activated processes are synapse and neuron projection, as well as hormones and amines secretion, due to the overexpression of several transcripts coding for synaptotagmins that act as sensors for calcium release. Among these transcripts, there were 35 lncRNAs (Fig. 1g), 20% of them being anti-sense to DNA-binding transcription factors (NR2F1, MEF2C, MYCBP2), calcium-dependent (CACNA2D1, SNAP25, SLC25A25), and RNA splicing (ZRANB2) related genes. Other lncRNAs include the brain-specific DLGAP2, known to be involved in synapse function and signaling in neuronal cells ^47^, or lncRNAs still not associated to neuronal tissue or diseases, like MHENCR, HAGLR, FAM225A and CEROX1.

Pseudobulk analyses strongly suggest that common mechanisms of transcriptional dysregulation can occur in ALS, as reflected by the many genes that are commonly de-regulated between the distinct glial cell populations (Supplementary Fig. 1e). Indeed, several differentially expressed lncRNAs were found in common between the cell subtypes, which include MZF1-AS1, shared between microglia and astrocytes, ZRANB2-AS1, shared between astrocytes and oligodendrocytes, and NR2F1-AS1, MYCBP2-AS1 and LINC01378 shared between microglia and oligodendrocytes (Fig. 1g). Interestingly, LINC01378 was previously shown linked by pleiotropy to schizophrenia and bipolar disorder ^48^. Pairwise correlations considering common DEGs between neuronal populations (Excitatory Neurons and Inhibitory Neurons) showed a quite perfect correlation (R squared = 0.9428, Supplementary Fig. 1 f), with three lncRNAs being up-regulated and 1 down-regulated. This could indicate common mechanisms of transcriptional activity between the two neuronal populations. Despite some lncRNAs were found dysregulated in two or more glial population, a lower correlation exists when considering all DEGs in glial cells, which could suggest different transcriptional mechanisms regulating cellular processes in those cell types.

Finally, in Oligodendrocyte Progenitor Cells (OPCs) pseudobulk analysis revealed 168 DEGs, the majority of which (70.5%) being up-regulated in ALS. GSEA analysis revealed that those transcripts are associated with the activation of GO terms mostly related to acids transports and endosomes, among others, and the suppression of RNA catabolic processes and unfolded protein response (Supplementary Table 2). Among the differentially expressed transcripts, 14 are annotated as lncRNAs, with 3 of them down-regulated and 11 up-regulated. 9 of the up-regulated lncRNAs were anti-sense RNAs, with none of their sense counterparts being differentially expressed. Among the down-regulated lncRNAs, we found CEROX1, a post-transcriptional regulator of mitochondrial activity ^49^, and MEG3, also found in previous ALS-linked bulk RNAseq data ^50^.

Taken together, these results indicate a possible role of lncRNAs in modulating the activation or suppression of distinct molecular mechanisms in glial cells of ALS patients, participating in the dysregulation of RNA processing, oxidative stress, calcium-driven processes, and synaptic organization.

### Dysregulation of lncRNAs in ALS neuronal populations

We next analyzed the neuronal populations, which include excitatory and inhibitory neurons. Excitatory neuron markers included 150 lncRNAs, 65 of which were anti-sense and 53 intergenic long non-coding RNAs (LINCs). Pseudobulk analysis revealed 78 lncRNAs that were differentially expressed between ALS and PN, with only 15 being down-sregulated in ALS. Seven of the differentially expressed lncRNAs between ALS and PN (LINC01106, MIR31HG, GRM5-AS1, CCDC144NL, PPFIA2-AS1, ZNF337-AS1 and DPYD-AS1) were also found as general excitatory neuron markers (computed as differentially expressed between excitatory neurons and all the other populations regardless the disease status), indicating a possible nature of these molecules as specific biomarkers of ALS excitatory neurons. LINC001106 is a lncRNAs whose expression was found high in bladder cancer patients ^51^ and in colorectal cancer cells ^52^, where it was found to be able to sponge miR-449b-5p and recruit FUS to GLI1 and GLI2 promoters to activate their transcription. In the snRNAseq dataset, we found it slightly down-regulated in ALS patients (log2FC=-0.58), which could imply a different mechanism of action of this lncRNA in this cell population. MIR31HG is a miRNA host gene that is up-regulated in BRAF-mutated melanoma cells and that can repress p16/CDKN2A expression by recruiting Polycomb group complexes ^53^. Its expression has been also positively correlated to invasiveness of colorectal cancer cells ^54^. However, in this ALS dataset, MIR31HG resulted slightly down-regulated (log2FC=-0.43).

Another candidate is GRM5-AS1, a still uncharacterized lncRNA found up-regulated (log2FC=1.44) in ALS and anti-sense to GRM5, a metabotropic glutamate receptor involved in neuronal network activity and synaptic plasticity. In ALS, glutamate receptors contribute to excitotoxicity and, in particular, the knock-down of GRM5 was shown to slow down disease progression in SOD mice ^55^. A high expression of GRM5 in ALS was also found in Positron Emission Tomography (PET) studies on ALS SOD mice ^56^. In such a situation, we could speculate that the high expression levels GRM5-AS1 in ALS patients is meant to counteract the neurotoxicity given by the hyperactivation of GRM5 receptors, but the specific mechanism of action of this lncRNA needs to be further studied. Other differentially expressed lncRNAs include PPFIA2-AS1. So far, no report has defined its role, but its sense transcript encode Liprin, a protein expressed in the brain and involved in the recruitment of dense core vesicles to post-synaptic sites ^57^. DPYD-AS1 is a lncRNA found up-regulated (log2FC=1.21) in ALS patients and anti-sense to DPYD, a gene coding for the Dihydropyrimidine Dehydrogenase enzyme. Despite no reports have shown a role for this lncRNA, deficiencies of the enzyme coded by the sense transcript could cause severe neurological manifestations in pediatric patients, as a consequence of the accumulation of thymine and uracil ^58^.

GSEA analysis revealed the suppression of processes related to neuron projection and cell cycle, while activated terms include G protein−coupled purinergic nucleotide receptor and DNA packaging (Supplementary Table 2). This is in line with what already observed in ALS pathogenesis, characterized primarily by the degeneration of motor neurons in which an impairment in neuronal projections could imply a disturbance of the anterograde and retrograde transport across axons. Purinergic signaling activation was observed to be beneficial in SOD1 mouse models ^59^, but we found purinergic receptors up-regulated in ALS patients. Such up-regulation could result from the glia hyperactivation and may suggest a possible mechanism of aberrant communications between the glial population and neurons in ALS.

In inhibitory neurons, we found 66 lncRNAs as positive or negative markers, with the majority of them (55) being positive markers. Pseudobulk analysis revealed only 10 lncRNAs that were differentially expressed between ALS and PN, none of which were also biomarkers for this cell population. Among them, were annotated as anti-sense RNAs, with their sense counterparts implicated in different annotated Gene Ontology terms, including glucocorticoids pathway, lipid metabolism and transcription factor activity regulation. GSEA analysis shows the enrichment of general cellular processes, including the suppression of growth factors signaling and cell death.

Interestingly, when looking at Genome Wide Association Studies for lncRNAs variants associated to ALS (MONDO_0004976 and EFO_0001357 from NHGRI-EBI Catalog ^60^), we found two lncRNAs that were also differentially expressed in ALS vs PN neuronal population. In particular SMIM2-AS1 (rs9533799) and LINC02742 (rs1491818) resulted up-regulated in excitatory neurons. Another lncRNA found in GWAS traits, KCNMB2-AS1 (rs13100616), was up-regulated in microglial population. All of these lncRNAs are poorly characterized, and further studies are needed to explore their possible involvement in ALS pathogenesis.

Taken together, these results point to a specific neuronal population, in particular excitatory neurons, as recipient of the distinct dysregulated processes observed in ALS over the past years and suggest further cellular processes that resulted impaired in ALS excitatory neurons. Moreover, it is suggested that distinct anti-sense RNAs could play a role in excitatory neurons of ALS patients, by participating in the activation or suppression of both general and neuronal-specific cell processes.

### Involvement of lncRNAs in the gene regulatory networks of ALS cell populations

To gain insights into the regulatory mechanisms underpinning ALS condition that cannot be captured by merely measuring transcript expression, we took advantage of the high dimensional Weighted Gene Co-expression Network analysis (hdWGCNA) ^61^. Specifically, we performed a consensus analysis to check the existence of gene regulatory networks (named modules) that resulted conserved between ALS and PN cell populations, and then used this analysis to find whether and to what extent those modules were dysregulated in ALS. The analysis revealed the presence of five distinct modules (Fig. 2 a), the majority of which being down-regulated in ALS, with the exception of the “turquoise” module that resulted significantly up-regulated (Fig. 2 b).

**Fig. 2.**
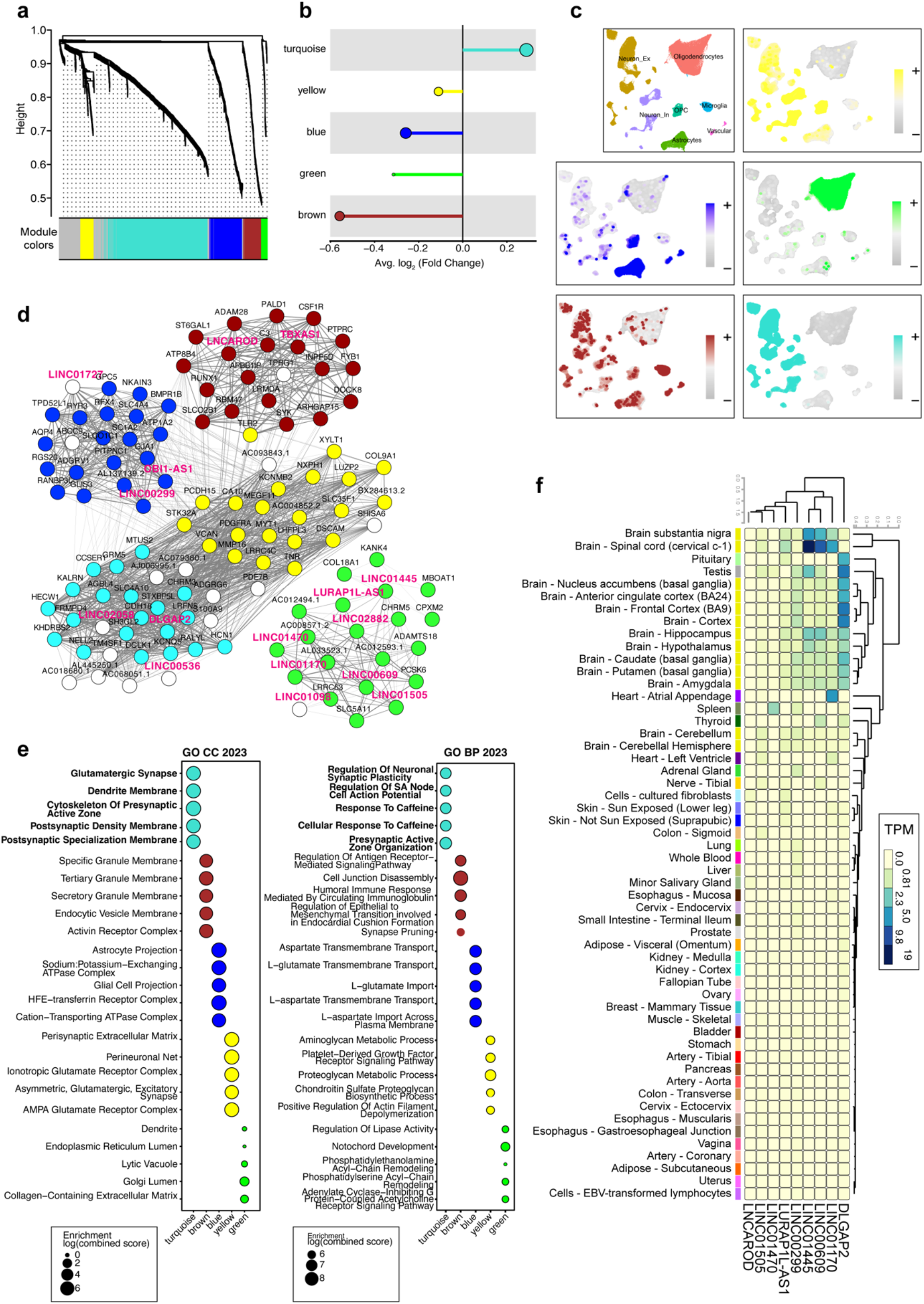
hdWGCNA analysis of cell populations. (a) Dendrogram derived from the consensus analysis (ALS vs PN) showing the existence of distinct color-coded modules. (b) Lollipop plot describing the up- and down-regulation of each module in ALS samples. (c) UMAP projections colored according to the kME values, that allow to assign each module to distinct cell populations. (d) Network of the top 20 hub genes derived from the consensus analysis, in which nodes are colored according to the module they belong to. White nodes are not assigned to a specific module or are not among the top 20 hub genes for any modules. LncRNA names are in red. (e) GSEA analysis of top 5 GO biological processes and GO Cellular components for each module. (f) Heatmap from GTEx resource showing the expression levels (TPM) of selected lncRNAs in the different tissues

We found that the turquoise module characterizes both excitatory ad inhibitory neuron populations (Fig. 2 c, d) and was enriched in genes belonging to glutamate receptor pathway and synaptic plasticity ontologies, in line with the importance of these processes in neuronal populations of ALS patients (Fig. 2 e). Conversely, the “blue” module was found enriched in astrocytes, it is down-regulated in ALS and its genes were enriched for processes related to aminoacidic and ion transport across membranes. Brown module enriched the microglia cell population, and its genes participate to processes related to activin-receptor and transforming growth factor beta signaling pathways, which resulted suppressed in ALS. Yellow module is active in neuronal population, particularly in inhibitory neurons, and its genes belong to cell components related to post synaptic density membrane, which resulted down-regulated in ALS. Finally, the “green” module enriched the oligodendrocyte population, it was found down-regulated in ALS and it is enriched in processes related to organelle organization and calcium ion channel activity, among others. Interestingly, 14 out of the top 100 hub genes of the five modules (top 20 for each module) are lncRNAs (Fig. 2 f). The majority of them belong to the “green” module, thus suggesting their possible role in the dysregulated mechanisms found in ALS oligodendrocytes. According to GTEx, 3 of these lncRNA, LINC01170, LINC00609 and LINC01445 are highly expressed in brain tissues (Fig. 2 g), with the higher expression in brain spinal cord. Among these lncRNAs, only one - LINC01445 – has been previously studied, as it has been found fused to EGFR in 17.7% of adult IDH-wt glioblastoma ^62^. Interestingly, the green module resulted unconnected to the rest of the network: as network edges represent co-expression relationships between genes, such a scenario indicates possible features (e.g. dysregulated cellular and molecular processes) which distinguish the green from the other modules.

Taken together, these results show how ALS transcriptome profiles are characterized by the down-regulation of distinct gene regulatory networks. As the 45% of the top hub genes of the “green” module include lncRNAs, we speculate a possible important contribution of lncRNAs to still unknown molecular processes that are altered in ALS astrocytes.

### No significant differences between C9orf72-mutated ALS and sporadic ALS

As the patient’s cohort included both C9orf72-mutated patients (n=6) and sporadic ALS patients (n=17), we took advantage of pseudobulk analysis to check differences between these two ALS subtypes. For each annotated cell type, no significant difference was found between the two ALS groups, except for two genes which were differentially expressed in almost all pseudobulk analyses. In particular, the still uncharacterized lncRNA C5orf17 (LINC02899) was strongly down-regulated in sALS *vs* c9ALS patients while the DEAD/H-Box Helicase DDX11 was slightly up-regulated.

Such a result is in line with a previous report were c9ALS data were found indistinguishable to sALS data by means of transcriptomics analysis from spinal cord samples ^36^, yet in contrast with other reports of analysis of frontal cortex and cerebellum regions from ALS and control patients ^63,64^. Due to the relatively low number of patients, real differences between the two ALS types should be further investigated with the use of increased sample sizes.

## Discussion

ALS is a neurodegenerative disease with unfavorable outcomes. Despite numerous efforts have been undertaken to define the primary mechanisms underlying the pathology, the purely genetic information that predisposes patients to disease remains insufficient, thus suggesting the contribution of additional processes. The lack of ideal model systems that perfectly emulate the human condition, coupled with challenges in obtaining cellular and tissue material for scientific analysis, further complicate these studies making research on ALS a non-trivial task.

To date, numerous cellular and molecular processes have been identified as dysregulated in ALS, primarily facilitated by the spread of OMICs technologies at both bulk and single-cell levels with a focus on distinct cellular populations within the brain and neuromuscular system. This has enabled a detailed exploration of molecular dynamics occurring in ALS cells, progressively adding more pieces to this puzzle. Some studies have focused on the regulation, processing, and metabolism of RNA, given the evident role it plays in transmitting genetic information from the genome to proteins. Particularly, RNA-associated processes have been observed to be dysregulated in ALS. Within this context, lncRNAs, a recently identified class of RNA that plays a crucial role in gene expression regulation, have emerged as particularly intriguing.

In numerous reports based on RNA sequencing data analyses, several lncRNAs have been found among differentially expressed transcripts, being either up- or down-regulated. However, such data are often derived from bulk experiments, which hinder a rapid and direct assignment of lncRNAs to specific cell types or subtypes, which is important due to their high cellular and tissue specificity. Furthermore, the significant lack of functional annotation for the majority of known lncRNAs greatly limits the ability to attribute a function to these molecules, including their involvement in crucial molecular and cellular processes. All of this could potentially introduce confounding variables, even at a statistical level, especially in cases where relatively high percentages of differentially expressed lncRNAs result from NGS experiments. In this study, we analyzed single-nuclei RNA sequencing data from one of the very few publicly available datasets, focusing on lncRNA expression within individual cellular populations. The more traditional single-cell analysis was integrated with tests of differential gene expression (pseudobulk) and network analyses. This led to the identification of a series of lncRNAs, the majority of which were not annotated, and that could be markers of neuronal populations. Moreover, some of these lncRNAs were found to be differentially expressed in glial or neuronal subpopulations, suggesting their potential being ALS-specific cell markers. Further comparison of gene regulatory networks in ALS patients and non-pathological controls led to the identification of several modules dysregulated in ALS. The genes belonging to those modules were then used for functional enrichment analyses, revealing how genes from each module relate to distinct deregulated biological processes in ALS. Moreover, one module, the green one, was particularly enriched in lncRNAs, indicating their likely prominent role within the specific cellular functions associated to that subnetwork. One limitation of this study is intrinsic in the type of single cell data which is restricted to the nuclear compartment and to the fraction of polyadenylated lncRNAs, which limits the study to those that can be captured by polyA-enriched methodologies. Overall, this analysis can show potential insights to generate new data-driven hypotheses, hence providing fertile ground for future cellular and molecular investigations.

## Methods

### Single nuclei RNA sequencing data processing

Single cell analysis has been performed on GEO datasets (accession number GSE174332) using Seurat package in R environment. Cells with less than 200 transcripts and with a percent of mitochondrial transcripts higher than 5% were filtered out. we randomly sub-sampled the dataset to 150,000 cell in total and followed a standard pipeline (log10 normalization, top 2000 variable features, data scaling, principal component analysis). We then used the first 30 principal components to find neighbors. Clusters were found using a resolution of 0.5. Markers were computed using the FindAllMarkers function in Seurat, retrieving both positive and negative markers, with the following parameters: min.pct=0.25, logfc.threshold=0.25, max.cells.per.ident=5000.

### Cell type identification in single cell RNA-seq analysis

For cell type identification we took advantage of both the already provided cell population annotation provided with the dataset, enriched with the expression of distinct known cell population markers that have been found by literature manual curation: SNAP25, SYT1, SLC17A7, SATB2, RORB, NEUROD2 for excitatory neurons, GAD1, GAD2, NXPH1 for inhibitory neurons, GFAP, AQP4, SLC1A2 for astrocytes, CSF1R, CD74, P2RY12, PTPRC, TMEM119 for microglia, MOBP, MBP, ST18, KLK6, SLC5A11 for oligodendrocytes, PDGFRA, CSPG4 for OPC, and ICAM1, PECAM1, FLT1 for vascular cells.

### Pseudobulk analysis

Pseudobulk analysis has been performed separately for each cell type by sub-setting the Seurat object for each specific cell population, and then aggregating data by summing the transcripts from all cells, obtaining a matrix with genes as rows and patients as columns. Differential expression analysis has been performed with DESeq2 ^65^ using the standard pipeline provided in the DESeq2 vignette. LncRNAs were downloaded from GENECODE database Release 44 (GRCh38.p14).

Enrichment analysis on differentially expressed genes has been performed using clusterProfiler R package ^66^ with the following parameters: nPerm=10000, minGSSize=3, maxGSSize=2000, pvalueCutoff=0.05). Heatmaps were drawn using pheatmap package. GTEx heatmaps were obtained on GTEx portal (https://www.gtexportal.org/home/) with the multi-gene query function. Whenever the number of lncRNA was higher than the maximum permitted by GTEx portal, we subset the number of lncRNAs to 100 per run.

### hdWGCNA analysis

hdWGCNA (high dimensional weighted gene co-expression network analysis) package ^61^ was used to identify gene co-expression networks of cells via unsupervised clustering. A consensus analysis between ALS and PN patients (n=23 and n=17, respectively) was performed using the hdWGCNA package in R environment according to the reference vignette of consensus analysis, changing some specific parameters as follows. In the “MetacellsByGroups function we set k=25, max_shared=12, min_cells=100, target_metacells=250. Soft power thresholds: 6 and 7. Gene ontology enrichment for each module and plotting was performed using EnrichR ^67^. The integrated network of all modules were retrieved with the top 20 hub genes from the hdWGCNA analysis as a table and then we used Cytoscape tool (v 3.7.1) ^68^ to draw the network and edit styles.

### Statistical analysis and figures

Statistical values in charts are reported as mean ± SEM unless otherwise specified. The statistical significance is defined as Bonferroni adjusted p-value where *p < 0.05; **p < 0.01; ***p < 0.001. For each type of bioinformatic analysis see the specific Materials and Methods section and Fig. captions. Figures were prepared and edited using Adobe Illustrator 2024 and Prism 7 (GraphPad) software.

## Data availability

The dataset used during the current study is available at the Gene Expression Omnibus (GEO) repository under the accession number GSE174332 (https://www.ncbi.nlm.nih.gov/geo/query/acc.cgi?acc=GSE174332). The code used for the analysis is available from the corresponding author, upon reasonable request.

## Acknowledgments

This work was supported by grants from European Union - Next-GenerationEU - National Recovery and Resilience Plan (NRRP) – MISSION 4 COMPONENT 2, INVESTMENT N. 1.1, CALL PRIN 2022 PNRR D.D.1409 of 14th Sep 2022 - P2022FFEWN, CUP: B53D23026140001); European Union - Next-GenerationEU - National Recovery and Resilience Plan (NRRP) – MISSION 4 COMPONENT 2, INVESTMENT N. 1.1, CALL PRIN 2022 - 2022BYB33L, CUP: B53D23016100006); European Union - NextGenerationEU: National Center for Gene Therapy and Drug based on RNA Technology, CN3 - Spoke 3 (code: CN00000041; PNRR MUR – M4C2 – Action 1.4-Call “Potenziamento strutture di ricerca e di campioni nazionali di R&S”, CUP: B83C22002870006) and Sapienza University (RM12117A5DE7A45B) to MB. AP salary is supported by the European Union - NextGenerationEU: National Center for Gene Therapy and Drug based on RNA Technology, CN3 - Spoke 3 (code: CN00000041; PNRR MUR – M4C2 – Action 1.4-Call “Potenziamento strutture di ricerca e di campioni nazionali di R&S”, CUP: B83C22002870006).

## Author contributions

CRediT authorship contribution: Conceptualization: AP, MB. Data curation, formal analysis, investigation and methodology: AP. Writing – review & editing: AP, MB. Project administration: AP, MB. Funding acquisition. MB. All authors have commented on, revised and approved the final version of the manuscript.

## Additional Information

The authors declare no competing interests.

---

**Supplementary Fig. 1.**
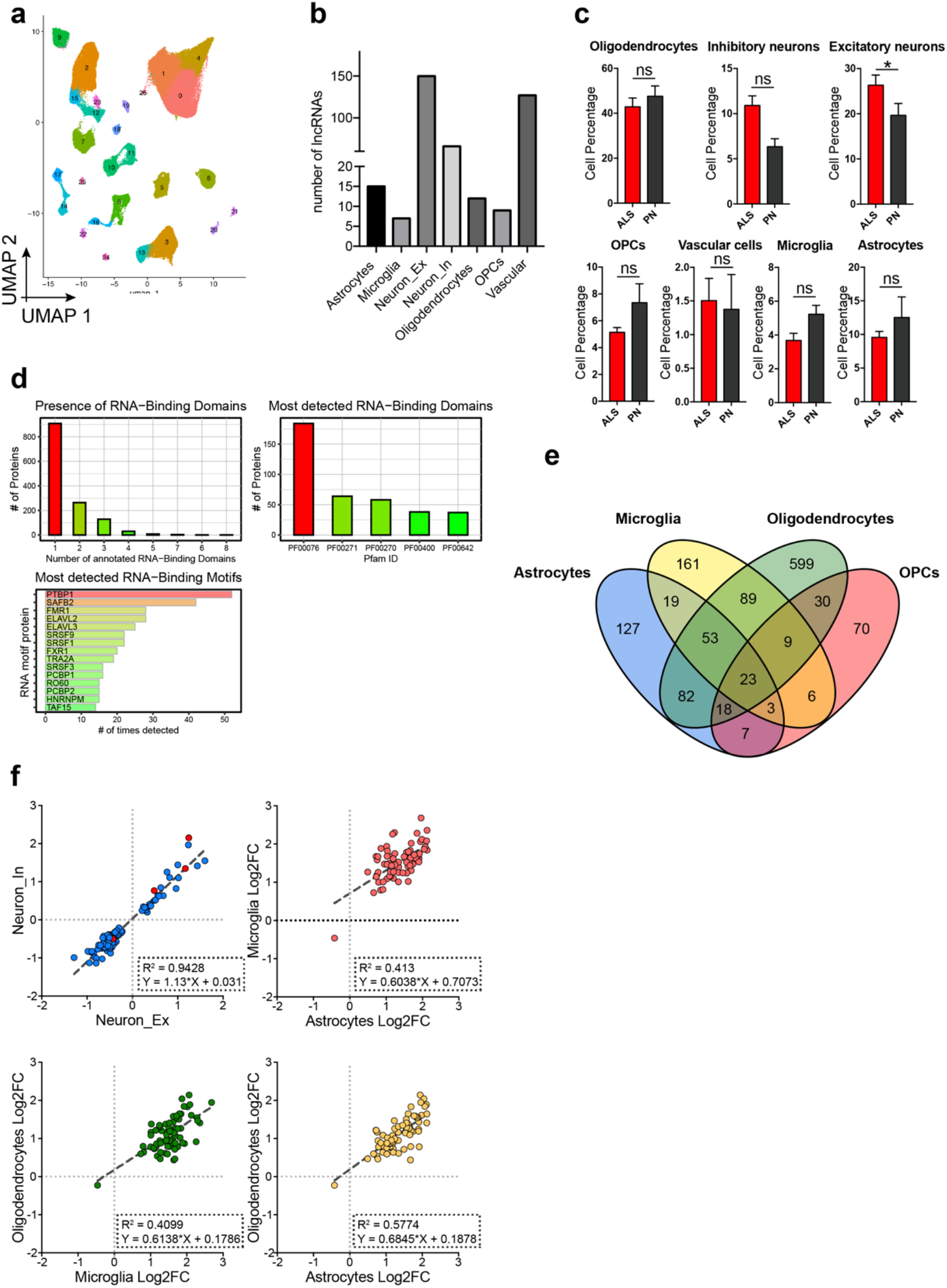
Analysis of cell populations and lncRNAs dynamics. (a) UMAP projection showing the clusters derived from the processing of the snRNAseq dataset. (b) Barplot reporting the number of lncRNAs that are differentially expressed in each cell population (ALS *vs* PN). (c) Percentage of cells belonging to each population in ALS and PN samples. (d) CatRAPID output for the BASP1-A1 transcript showing its protein-binding predictions. (e) Venn diagram of differentially expressed lncRNAs in the distinct cell populations. (f) Expression correlation plots (log2 fold change) of lncRNA divided by neuronal populations and glial populations. For glial populations, only lncRNAs in common between all three cell populations are shown.

## References

1. Hardiman, O. et al. Amyotrophic lateral sclerosis. Nature Reviews Disease Primers 3, 17071 (2017).

2. Xu, L. et al. Global variation in prevalence and incidence of amyotrophic lateral sclerosis: a systematic review and meta-analysis. J. Neurol. 267, 944–953 (2020).

3. Sau, D. et al. Mutation of SOD1 in ALS: A gain of a loss of function. Hum. Mol. Genet. 16, 1604–1618 (2007).

4. DeJesus-Hernandez, M. et al. Expanded GGGGCC Hexanucleotide Repeat in Noncoding Region of C9ORF72 Causes Chromosome 9p-Linked FTD and ALS. Neuron 72, 245–256 (2011).

5. Renton, A. E. et al. A hexanucleotide repeat expansion in C9ORF72 is the cause of chromosome 9p21-linked ALS-FTD. Neuron 72, 257–268 (2011).

6. Van Deerlin, V. M. et al. TARDBP mutations in amyotrophic lateral sclerosis with TDP-43 neuropathology: a genetic and histopathological analysis. Lancet Neurol. 7, 409–416 (2008).

7. Jr, K. T. J. et al. Mutations in the FUS/TLS gene on chromosome 16 cause familial amyotrophic lateral sclerosis. Science (80-. ). 323, 1205–1208 (2009).

8. Maruyama, H. et al. Mutations of optineurin in amyotrophic lateral sclerosis. 465, 223–226 (2010).

9. Fecto, F. et al. SQSTM1 mutations in familial and sporadic amyotrophic lateral sclerosis. Arch. Neurol. 68, 1440–1446 (2011).

10. Ho, W. Y. et al. The ALS-FTD-linked gene product, C9orf72, regulates neuronal morphogenesis via autophagy. Autophagy 15, 827–842 (2019).

11. Yang, M. et al. A C9ORF72/SMCR8-containing complex regulates ULK1 and plays a dual role in autophagy. Sci. Adv. 2, (2016).

12. Kok, J. R., Palminha, N. M., Dos Santos Souza, C., El-Khamisy, S. F. & Ferraiuolo, L. DNA damage as a mechanism of neurodegeneration in ALS and a contributor to astrocyte toxicity. Cell. Mol. Life Sci. 78, 5707–5729 (2021).

13. Akçimen, F. et al. Amyotrophic lateral sclerosis: translating genetic discoveries into therapies. Nat. Rev. Genet. (2023). doi:10.1038/S41576-023-00592-Y

14. Thuault, S. The RNAs of ALS. Nature Neuroscience 18, 1066 (2015).

15. Lee, Y. J. & Rio, D. C. A mutation in the low-complexity domain of splicing factor hnRNPA1 linked to amyotrophic lateral sclerosis disrupts distinct neuronal RNA splicing networks. Genes Dev. 38, 11–30 (2024).

16. Verdile, V. et al. Dysregulation of alternative splicing underlies synaptic defects in familial amyotrophic lateral sclerosis. Prog. Neurobiol. 231, (2023).

17. Ramaswami, M., Taylor, J. P. & Parker, R. Altered ribostasis: RNA-protein granules in degenerative disorders. Cell 154, (2013).

18. Raguseo, F. et al. The ALS/FTD-related C9orf72 hexanucleotide repeat expansion forms RNA condensates through multimolecular G-quadruplexes. Nat. Commun. 14, (2023).

19. Naskar, A., Nayak, A., Salaikumaran, M. R., Vishal, S. S. & Gopal, P. P. Phase separation and pathologic transitions of RNP condensates in neurons: implications for amyotrophic lateral sclerosis, frontotemporal dementia and other neurodegenerative disorders. Front. Mol. Neurosci. 16, 1–18 (2023).

20. Ziff, O. J. et al. Nucleocytoplasmic mRNA redistribution accompanies RNA binding protein mislocalization in ALS motor neurons and is restored by VCP ATPase inhibition. Neuron 111, 3011–3027.e7 (2023).

21. Rajabi, D., Khanmohammadi, S. & Rezaei, N. The role of long noncoding RNAs in amyotrophic lateral sclerosis. Rev. Neurosci. (2024). doi: 10.1515/revneuro-2023-0155

22. Laneve, P., Tollis, P. & Caffarelli, E. Rna deregulation in amyotrophic lateral sclerosis: The noncoding perspective. Int. J. Mol. Sci. 22, 1–34 (2021).

23. Ballarino, M., Morlando, M., Fatica, A. & Bozzoni, I. Non-coding RNAs in muscle differentiation and musculoskeletal disease. J. Clin. Invest. 126, 2021–2030 (2016).

24. Buonaiuto, G., Desideri, F., Taliani, V. & Ballarino, M. Muscle Regeneration and RNA: New Perspectives for Ancient Molecules. 1–21 (2021).

25. Pagano, F., Calicchio, A., Picchio, V. & Ballarino, M. The Noncoding Side of Cardiac Differentiation and Regeneration. Curr. Stem Cell Res. Ther. 15, 723–738 (2020).

26. Rosa, A. & Ballarino, M. Long Noncoding RNA Regulation of Pluripotency. Stem Cells Int. 2016, (2016).

27. Mattick, J. S. et al. Long non-coding RNAs: definitions, functions, challenges and recommendations. Nat. Rev. Mol. Cell Biol. 1–17 (2023). doi:10.1038/s41580-022-00566-8

28. Statello, L., Guo, C. J., Chen, L. L. & Huarte, M. Gene regulation by long non-coding RNAs and its biological functions. Nature Reviews Molecular Cell Biology 22, 96–118 (2021).

29. Ferrè, F., Colantoni, A. & Helmer-Citterich, M. Revealing protein-lncRNA interaction. Brief. Bioinform. 17, 106–116 (2016).

30. Ribeiro, D. M. et al. Protein complex scaffolding predicted as a prevalent function of long non-coding RNAs. Nucleic Acids Res. 46, 917–928 (2018).

31. Kretz, M. et al. Control of somatic tissue differentiation by the long non-coding RNA TINCR. Nature 493, 231–235 (2013).

32. Carvelli, A. et al. A multifunctional locus controls motor neuron differentiation through short and long noncoding RNAs. EMBO J. 41, e108918 (2022).

33. Ballarino, M., Pepe, G., Helmer-Citterich, M. & Palma, A. Exploring the landscape of tools and resources for the analysis of long non-coding RNAs. Comput. Struct. Biotechnol. J. 21, 4706–4716 (2023).

34. Gagliardi, S. et al. Long non coding RNAs and ALS: Still much to do. Non-coding RNA Res. 3, 226–231 (2018).

35. Aliperti, V., Skonieczna, J. & Cerase, A. Long non-coding rna (Lncrna) roles in cell biology, neurodevelopment and neurological disorders. Non-coding RNA 7, (2021).

36. Humphrey, J. et al. Integrative transcriptomic analysis of the amyotrophic lateral sclerosis spinal cord implicates glial activation and suggests new risk genes. Nat. Neurosci. 26, 150–162 (2023).

37. Pineda, S. S. et al. Single-cell profiling of the human primary motor cortex in ALS and FTLD. bioRxiv 2021.07.07.451374 (2021). doi:10.1101/2021.07.07.451374

38. Lonsdale, J. et al. The Genotype-Tissue Expression (GTEx) project. Nat. Genet. 45, 580–585 (2013).

39. Guttenplan, K. A. et al. Knockout of reactive astrocyte activating factors slows disease progression in an ALS mouse model. Nat. Commun. 11, 1–9 (2020).

40. Haidet-Phillips, A. M. et al. Astrocytes from familial and sporadic ALS patients are toxic to motor neurons. Nat. Biotechnol. 29, 824–828 (2011).

41. Nagai, M. et al. Astrocytes expressing ALS-linked mutated SOD1 release factors selectively toxic to motor neurons. Nat. Neurosci. 10, 615–622 (2007).

42. Andrés-Benito, P., Moreno, J., Aso, E., Povedano, M. & Ferrer, I. Amyotrophic lateral sclerosis, gene deregulation in the anterior horn of the spinal cord and frontal cortex area 8: Implications in frontotemporal lobar degeneration. Aging (Albany. NY). 9, 823–851 (2017).

43. Armaos, A., Colantoni, A., Proietti, G., Rupert, J. & Tartaglia, G. G. CatRAPID omics v2.0: Going deeper and wider in the prediction of protein-RNA interactions. Nucleic Acids Res. 49, W72–W79 (2021).

44. Mori, K. et al. HnRNP A3 binds to GGGGCC repeats and is a constituent of p62-positive/TDP43-negative inclusions in the hippocampus of patients with C9orf72 mutations. Acta Neuropathol. 125, 413–423 (2013).

45. Obrador, E. et al. The link between oxidative stress, redox status, bioenergetics and mitochondria in the pathophysiology of als. Int. J. Mol. Sci. 22, (2021).

46. Kang, S. H. et al. Degeneration and impaired regeneration of gray matter oligodendrocytes in amyotrophic lateral sclerosis. Nat. Neurosci. 16, 571–579 (2013).

47. Ouellette, A. R. et al. Cross-Species Analyses Identify Dlgap2 as a Regulator of Age-Related Cognitive Decline and Alzheimer’s Dementia. Cell Rep. 32, 108091 (2020).

48. Polushina, T. et al. Identification of pleiotropy at the gene level between psychiatric disorders and related traits. Transl. Psychiatry 11, (2021).

49. Sirey, T. M. et al. The long non-coding rna cerox1 is a post transcriptional regulator of mitochondrial complex i catalytic activity. Elife 8, 1–31 (2019).

50. Ruffo, P. et al. Deregulation of ncRNA in Neurodegenerative Disease: Focus on circRNA, lncRNA and miRNA in Amyotrophic Lateral Sclerosis. Front. Genet. 12, 1–10 (2021).

51. Meng, L., Xing, Z., Guo, Z. & Liu, Z. LINC01106 post-transcriptionally regulates ELK3 and HOXD8 to promote bladder cancer progression. Cell Death Dis. 11, 1–15 (2020).

52. Guo, K. et al. LINC01106 drives colorectal cancer growth and stemness through a positive feedback loop to regulate the Gli family factors. Cell Death Dis. 11, 1–15 (2020).

53. Montes, M. et al. The long non-coding RNA MIR31HG regulates the senescence associated secretory phenotype. Nat. Commun. 12, 1–17 (2021).

54. Eide, P. W., Eilertsen, I. A., Sveen, A. & Lothe, R. A. Long noncoding RNA MIR31HG is a bona fide prognostic marker with colorectal cancer cell-intrinsic properties. Int. J. Cancer 144, 2843–2853 (2019).

55. Bonifacino, T. et al. In-vivo genetic ablation of metabotropic glutamate receptor type 5 slows down disease progression in the SOD1G93A mouse model of amyotrophic lateral sclerosis. Neurobiol. Dis. 129, 79–92 (2019).

56. Brownell, A. L. et al. PET imaging studies show enhanced expression of mGluR5 and inflammatory response during progressive degeneration in ALS mouse model expressing SOD1-G93A gene. J. Neuroinflammation 12, 217 (2015).

57. Stucchi, R. et al. Regulation of KIF1A-Driven Dense Core Vesicle Transport: Ca2+/CaM Controls DCV Binding and Liprin-α/TANC2 Recruits DCVs to Postsynaptic Sites. Cell Rep. 24, 685–700 (2018).

58. Maekawa, K. et al. Genetic variations and haplotype structures of the DPYD gene encoding dihydropyrimidine dehydrogenase in Japanese and their ethnic differences. J. Hum. Genet. 52, 804–819 (2007).

59. Andries, M., Van Damme, P., Robberecht, W. & Van Den Bosch, L. Ivermectin inhibits AMPA receptor-mediated excitotoxicity in cultured motor neurons and extends the life span of a transgenic mouse model of amyotrophic lateral sclerosis. Neurobiol. Dis. 25, 8–16 (2007).

60. Sollis, E. et al. The NHGRI-EBI GWAS Catalog: knowledgebase and deposition resource. Nucleic Acids Res. 51, D977–D985 (2023).

61. Morabito, S., Reese, F., Rahimzadeh, N., Miyoshi, E. & Swarup, V. hdWGCNA identifies co-expression networks in high-dimensional transcriptomics data. Cell Reports Methods 3, (2023).

62. Arora, S. et al. Visualizing genomic characteristics across an RNA-Seq based reference landscape of normal and neoplastic brain. Sci. Rep. 13, (2023).

63. Prudencio, M. et al. Distinct brain transcriptome profiles in C9orf72-associated and sporadic ALS. Nat. Neurosci. 18, 1175–1182 (2015).

64. Dickson, D. W. et al. Extensive transcriptomic study emphasizes importance of vesicular transport in C9orf72 expansion carriers. Acta Neuropathol. Commun. 7, 1–21 (2019).

65. Love, M. I., Huber, W. & Anders, S. Moderated estimation of fold change and dispersion for RNA-seq data with DESeq2. Genome Biol. 15, 550 (2014).

66. Wu, T. et al. clusterProfiler 4.0: A universal enrichment tool for interpreting omics data. Innovation 2, 100141 (2021).

67. Kuleshov, M. V. et al. Enrichr: a comprehensive gene set enrichment analysis web server 2016 update. Nucleic Acids Res. 44, W90 (2016).

68. Shannon, P. et al. Cytoscape: A Software Environment for Integrated Models of Biomolecular Interaction Networks. Genome Res. 13, 2498–2504 (2003).

